# Cell Cycle Asynchrony Generates DNA Damage at Mitotic Entry in Polyploid Cells

**DOI:** 10.1101/582460

**Authors:** Maddalena Nano, Anthony Simon, Carole Pennetier, Vincent Fraisier, Veronique Marthiens, Renata Basto

## Abstract

Polyploidy arises from the gain of complete chromosomes sets [1] and is known to promote cancer genome evolution. Recent evidence suggests that a large proportion of human tumours experience whole genome duplications (WGDs), which might favour the generation of highly abnormal karyotypes within a short time frame, rather than in a stepwise manner [2–6]. However, the molecular mechanisms linking whole genome duplication to genetic instability remain poorly understood. Further, possible mechanisms responsible for rapid genome reshuffling have not been described yet. Using repeated cytokinesis failure to induce polyploidization of *Drosophila* neural stem cells (NSCs, also called neuroblasts - NBs), we investigated the consequences of polyploidy *in vivo*. Here, we show that polyploid NSCs accumulate high levels of chromosome instability. Surprisingly, we found that DNA damage is generated in a subset of nuclei of polyploid NBs during mitosis, in an asymmetric manner. Importantly, our observations in flies were confirmed in mouse NSCs (mNSCs) after acute cytokinesis inhibition. Interestingly, DNA damage occurs in nuclei that were not ready to enter mitosis but were forced to do so when exposed to the mitotic environment of neighbouring nuclei within the same cell. Additionally, we found that polyploid cells are cell cycle asynchronous and forcing cell cycle synchronization is sufficient to lower the levels of DNA damage generated during mitosis. Overall, this work supports a model in which DNA damage at mitotic entry can generate a mutated genetic landscape that contributes to the onset of genetic instability.

## Results and Discussion

### *Drosophila* neural stem cells as a model to study the consequences of polyploidy *in vivo*

To identify the impact of polyploidization *in vivo*, we chose the neural stem cells (NSCs) (also called neuroblasts, NBs) of the developing larval brain of *Drosophila*, which can accumulate high number of chromosomes in certain mutant conditions [7–9]. Polyploidy can be induced through the use of mutations or RNA interference (RNAi) that target genes required for cytokinesis such as the non-muscle myosin II regulatory light chain, *spaghetti squash* (*sqh*), Pavarotti (*pav*) (MKLP-1) and Anillin (*anl*),[8, 10, 11]. Polyploid brains contained NBs of variable size that were in the large majority multinucleated (Figure 1A-B) and maintained NSC fate as illustrated by Deadpan expression, a NSC marker [12–14](Supplementary Figure 1A). Not all *Drosophila* proliferative tissues were tolerant to polyploidy. Analysis of mutant wing discs, revealed severe development and proliferative defects accompanied by the presence of pyknotic nuclei, consistent with cell death reported in this tissue after cytokinesis inhibition [15] (Supplementary Figure 1B and data not shown). Importantly, inducing polyploidy in the a wild type (WT) background, either is discs or brains using different means to induce cytokinesis failure, confirmed the results described above (Supplementary Figure 1C-D). Large polyploid NB clones were detected in the brain, while in the disc this was not the case. We concluded that in the developing brain, but not in wing discs, polyploid cells can be maintained regardless of the fitness of neighbouring cells.

**Figure 1.**
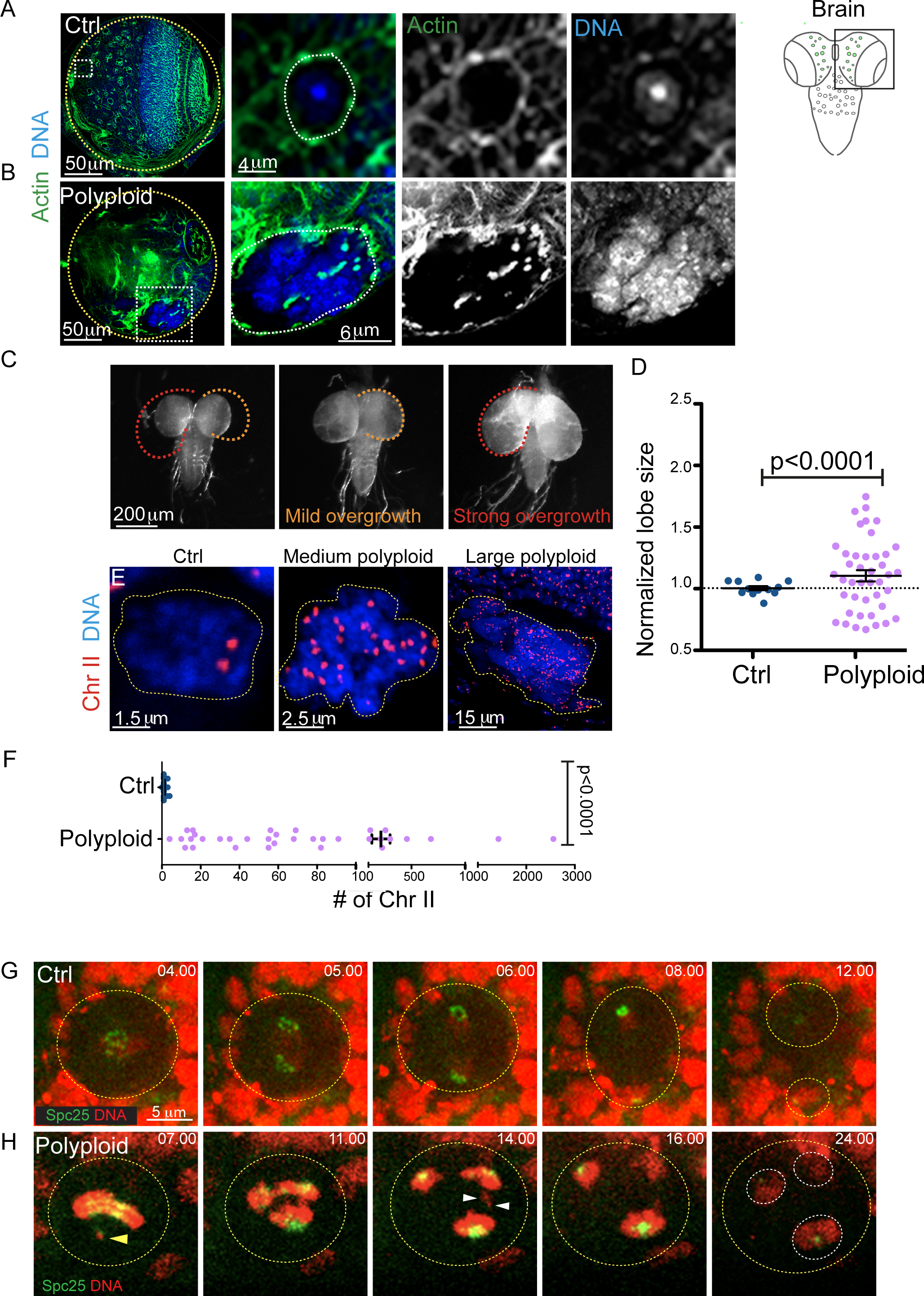
Polyploidy has tissue-specific consequences during *Drosophila* development and leads to genetic instability in NBs. (A-B) On the right, diagram showing a *Drosophila* 3^rd^ instar larval brain. Larval NBs are depicted in green. One of the two brain lobes is highlighted with a square. On the left, micrograph of diploid Ctrl (A) and polyploid (B) brain lobes labelled with Phalloidin to show F-actin (green) and the DNA is in blue. Yellow dashed lines highlight the boundary of the brain lobes, yellow dashed circles highlight NBs, while white squares indicate the region magnified in the inset on the right. (C) Ctrl (left) and polyploid (right) brains displaying mild or strong overgrowth. The orange dashed line outlines a polyploid brain lobe with mild overgrowth, while the red dashed line outlines a polyploid brain lobe with strong overgrowth. Dashed lines are superimposed to the Ctrl brain highlighting the difference in size. (D) Dot plot graph of normalized brain size in Ctrl (blue) and polyploid (pink) mutant brains Statistical significance of difference between variances was assessed by F-test. (E) Fluorescent insitu hybridization (FISH) with probes directed against chromosome II (Chr II, in red) of Ctrl, medium and large polyploid NBs. DNA is shown in blue. Yellow dashed lines highlight NB nuclear contours. (F) Dot plot chart showing the numbers of Chr II in Ctrl and polyploid NBs. Statistical significances were asses by an unpaired t-test. (G-H) Stills of time-lapse movies of mitotic NBs expressing H2Av fused to RFP to visualize DNA (shown in red) and Spc25 fused to GFP to visualize the kinetochore (shown in green). Yellow dashed lines outline cells boundaries, white dashed lines outline the daughter nuclei formed at the end of mitosis. Time is shown in minutes.seconds. Time zero was considered at NEBD. (G) Representative mitosis of a Ctrl NB. Note how all chromosomes in a given NB are associated to Spc25 signal (n=16 NBs). (H) Polyploid NB mitosis. At time 07.00 a chromosome with Spc25 signal still needs to align on the metaphase plate (yellow arrowhead). At anaphase (from t=11.00 to t=16.00), an acentric chromosome lacking Spc25 is noticed (white arrowheads, n=9/38NBs having chromosomes that lack Spc25). The acentric chromosome lags behind the main chromosome mass. Despite its delay, the acentric chromosome is ultimately segregated to opposite poles and incorporated in the newly formed nuclei.

We characterized in more detail the consequences of repetitive cytokinesis failure through the use of a *sqh* hypomorphic mutation [8]. Due to the hypomorphic nature of this allele, at late larval stages, mutant brains contain NBs with a highly variable degree of polyploidy (as some have failed cytokinesis only once or twice, while others have failed cytokinesis several times consecutively). For simplicity, we will refer to *sqh* mutants as polyploid throughout this work and we will call all diploid conditions used control (Ctrl; for complete genotypes and conditions please refer to figure legends and methods).

When compared to Ctrls, polyploid brains were highly variable in size as determined by lobe area (Ctrl: 0.71±0.02μm^2^, n=51 lobes and polyploid: 0.92±0.04, n=42, p<0.0001) (Figure 1C-D), with high proportion of brains showing strong overgrowth. Having established our system as a suitable model to study the evolution of polyploidization, we focused on the characterization of polyploid NBs.

### Polyploid NBs accumulate high number of incomplete chromosome sets

In order to determine the chromosomal content of polyploid NBs, we performed Fluorescence In Situ Hybridization (FISH) using probes for chromosome II. In Ctrl NBs, we could either detect one or two signals (Figure 1E-F), while in polyploid NBs the number ranged from four to 2559, which corresponds to roughly 10 rounds of cytokinesis failure (Figure 1E-F).

While characterising mitosis in polyploid NBs, we observed acentric chromosomes (Figure 1G-H). This was observed in brains expressing H2Av-RFP to visualize DNA and the kinetochore protein Spc25 (a component of the Ncd80 complex) [16]. Acentric chromosomes are characteristic of cells that experience genomic instability [17] and have been observed in *Drosophila* papillar cells [18]. In Ctrl cells we never observed mitotic chromosomes lacking Spc25 (n=16 NBs from 2 brains) (Figure 1G). However, polyploid cells often contained DNA fragments that lacked Spc25 signal (Figure 1H). Further, they lagged behind the main chromosome mass, separating and segregating to opposite poles only much later (n=9/38 cells from 8 brains), as described for acentric chromosomes induced by the I-CreI inducible system [19]. Importantly, acentric fragments are normally connected to the main chromatin mass by fine DNA threads that recruit proteins like BubR1 and Polo [19]. Using a FlyTrap line for GFP-Polo [20], we were able to visualize its transient recruitment to the thread linking acentric chromosomes to the main chromosome mass in polyploid NBs (11/11 NBs from 4 brains; Supplementary Figure 1F). This type of event was never detected in Ctrl NBs (Supplementary Figure 1E) [19]. These results suggest that DNA structural alterations-induced by DNA damage, which has been described occuring in polyploidy cells [5] was present in polyploid NBs and could contribute to chromosome instability typical of cells undergoing whole genome duplications (WGDs) [2, 3].

### DNA damage is detected in polyploid *Drosophila* NBs and mouse NSCs in mitosis

The observations of DNA structural alterations by time lapse microscopy in polyploid NBs prompted us to investigate the presence of DNA damage in NBs. We used the double strand break (DSB) marker γ–H2Av, which is one of earliest markers of the DNA damage response (DDR) [21]. As expected, we observed DNA damage in polyploid cells in interphase (data not shown). However, upon the analysis of polyploid brains, we noticed that certain mitotic cells contained γ–H2Av signals. Moreover, this signal seemed to be restricted to a certain area of the DNA and did not occupy the entire chromosome mass (Figure 2A-B). Since, to our knowledge, this type of defect has not been described before, and was never detected in Ctrl NBs (Figure 2A), we focused our analysis on these cells

**Figure 2.**
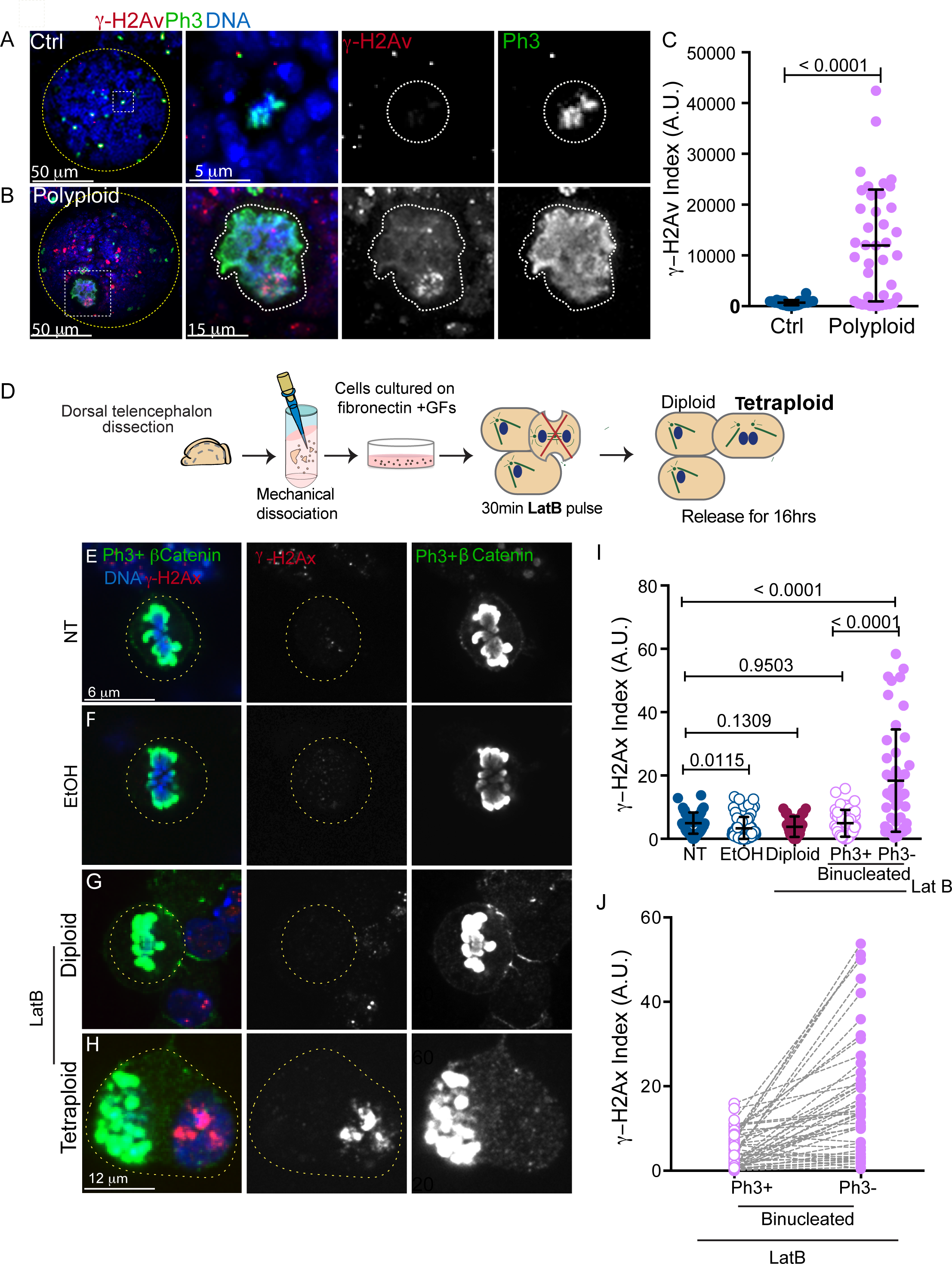
Mitotic polyploid NBs and mNSCs present DNA damage. (A-B) Micrographs of Ctrl and polyploid brain lobes (left) and mitotic NBs (right) stained with antibodies detecting DNA double-strand breaks (γ-H2Av, in red in the merged panel and in gray) and Ph3 (which labels mitotic cells, in green in the merged panel and in gray). DNA is shown in blue. The yellow dashed lines outlines the brain lobes, while the white dashed lines surround the NBs magnified in the insets. (C) Dot plot chart showing γ-H2Av index (expressed in arbitrary units – A.U.) measured in mitotic Ctrl (blue) and polyploid cells (pink). (D) Schematic representation of the workflow used to to establish and favour the propagation of pure primary adherent cultures of mouse embryonic neural stem cells (mNSCs). E16.5 cerebral cortex explants were enzymatically and mechanically dissociated into a single cell suspension, plated on fibronectin substrates and cultured in proliferation medium. A 30 min Latrunculin-B (LatB) treatment was used to induce cytokinesis failure; cells were then released for 16 hrs before being analyzed. (E-H) Micrographs of non-treated (E; NT) ethanol-treated (F; EtOH) and diploid (G) and tetraploid (H) Latrunculin-B-treated (LatB) mNSCs labelled with antibodies directed against γ-H2Ax to label DNA damage (in red in the merged panel and in gray) and Ph3 to label mitotic cells (in green in the merged panel and in gray). DNA is shown in blue. (I) Dot plot chart showing the γ-H2Ax index measured in mitotic mNSCs non-treated (NT, blue), ethanol-treated (EtOH, blue) and LatB-treated (pink). LatB-treated mNSCs were divided in sub-cathegories according to their ploidy (diploid, or binucleated). The two nuclei of LatB-treated, binucleated mNSCs are shown separately on the chart according to their mitotic status (Ph3+ and Ph3-). (J) “Before and after” chart highlighting the difference in γ-H2Ax index of mitotic (Ph3+) versus non mitotic nuclei (Ph3-) of asynchronous, binucleated mNSCs deriving from LatB treatment. Note how non-mitotic nuclei are more vulnerable to DNA damage. Statistical significance was assessed by unpaired t-test.

In Ctrl brains, γ–H2Av appears as a faint nuclear signal and only rare foci could be identified (Figure 2A). In mitotic polyploid NBs where signals for γ–H2Av were present, these were asymmetricly distributed and restricted to localized regions of the polyploid nucleus, while the rest of the DNA appeared to contain less γ–H2Av signal (Figure 2B). To accurately assess the frequency and intensity of DSBs, we applied a quantitative approach at the single cell level using a new plugin to measure both mean fluorescence intensity (FI) of γ–H2Av signals and the percentage of nuclear area covered by this signal (coverage, see methods). By multiplying the FI by the coverage, we obtained what we called the γ–H2Av index, which was highly increased in mitotic polyploid NBs compared to Ctrl NBs (Ctrl: 700.4± 83.2.01, n=36 NBs and polyploid: 11960±1743, n=40 NBs, p<0.0001) (Figure 2C).

In polyploid NBs, multinucleation is established by multipolar mitosis [22]. We wanted to assess wether a decrease in the extent of mitotic spindle multipolarity could reduce DNA damage in mitotic cells. To limit the number of mitotic spindle poles, we used the strategy described in [22], where a mutation in the centrosome encoding gene Sas-4 (Sas-4^mut^) [23] can be used to decrease the frequency of polyploid multipolar divisions. Importantly, Sas-4^mut^ mitotic NBs showed low levels of DNA damage, similar to Ctrls (Ctrl: 626.2.4±90.5, Sas-4^mut^: 672.3±167.4, p=0.7958; n=51 NBs and n=37 NBs respectively) (Supplementary Figure 2A and 2C). Even if a decrease in multipolarity resulted in a slight decrease in the γ–H2Av index in polyploid, Sas-4^mut^ mitotic cells, this was not significant (Supplementary Figure 2B-C) (Polyploid: 9183.0±2568, Polyploid, Sas-4^mut^: 7433±1149, p=0.4735, n=29 NBs and n=73 NBs respectively). Together, these results show that reducing multipolarity does not greatly influence the DNA damage found in polyploid mitotic cells.

We next tested whether mammalian polyploid cells could also present DNA damage in mitosis. By analogy to the system described above in flies, we established a mammalian model system where primary cultures of highly proliferative, dissociated mouse NSCs (mNSCs) can be established and maintained on fibronectin subtrates supplemented with growth factors [24] (Figure 2D). We established mNSCs primary cultures derived from E16.5 embryos. We used a 30 min Latrunculin-B (LatB) pulse to inhibit actin polymerisation, with the aim of inhibiting cytokinesis in cells that were in telophase (Figure 2D). LatB treatment was followed by washout and cells were allowed to recover for 16 hrs. 11.4% of LatB-treated cells (n=320 out of 2819 cells analysed) contained extra centrosomes, while this frequency was only 0.5% (n=8 cells out of 1668 cells analysed) in non treated (NT) mNSCs and 0.6% in ethanol-treated mNSCs (Lat B is diluted in ethanol) (n=10 out of 1749 cells analysed) (Supplementary Figure 2D-E). We restrained the analysis of these cultures to a period of 16hrs after cytokinesis inhibition, which represents roughly two cell cycles (our data), since we noticed a significant drop in proliferation afterwards (not shown), consistent with the presence of mechansims that eliminate unscheduled polyploidization in vertebrate cells [25, 26].

We next quantified γ-H2Ax fluorescent levels. Since the signals obtained were different from the signal found in *Drosophila* NBs, and more spread in volume, we decided to generate another plugin that takes into account γ-H2Ax signals through multiple image stacks. In mitotic NT, EtOH treated or even mononucleated LatB (presumably diploid) mNSCs, γ–H2Ax index was very low (NT: 5.0±0.45, n=54 cells, EtOH:3.4±0.4, n=66 cells and 3.8±0.5, n=29 cells) (Figure 2E-G and 2I). Strikingly, in a large proportion of binucleated cells, we could observed that the nucleus that contained higher γ–H2Ax levels was less advanced in mitosis (defined by Ph3 FI levels or by chromosome condensation) (Ph3+: 4.9±0.6 and Ph3-:18.4±2.4, n=47) (Figure 2G and 2I). Additionally, plotting γ–H2Ax levels in the “before and after” graph type, showed a considerable increase of DNA damage levels in the nucleus that contained lower Ph3 levels within the same cell (Figure 2J).

We concluded that both polyploid *Drosophila* NSCs and binucleated mNSCs present DNA damage in mitosis.

### DNA damage is generated during mitosis in polyploid NBs

The fact that certain nuclei/nuclear domains in polyploidy cells contained high levels of DNA damage suggested at least two possibilities. DNA damage could occur during mitosis in certain vulnerable nuclei or nuclear regions, but not in others. Alternatively, these nuclei or nuclear regions, could enter mitosis with DNA damage generated in the previous interphase. To distinguish between these possibilities, we used RpA-70-GFP [27] in time-lapse movies of *Drosophila* NBs. RpA-70 is a member of the elongation factor complex and binds single-stranded DNA (ssDNA) produced through resection after the generation of dsDNA breaks [28]. We first tested whether we could image Ctrl brains expressing RpA-GFP for extended periods of time without detecting the appearance of foci. This was indeed the case, as in movies over 300min (5hrs) NBs continued to divide without accumulating ssDNA (Figure 3A and Movie 1). We then verified the use of RpA as a marker for DNA damage in this experimental system. We induced DNA damage using laser ablation in diploid Ctrl NBs. RpA-70-GFP was rapidly localized to focal structures within the nucleus (Figure 3B and Movie 2), validating the use of this line and the efficiency of RpA recrutement upon induced DNA damage.

**Figure 3.**
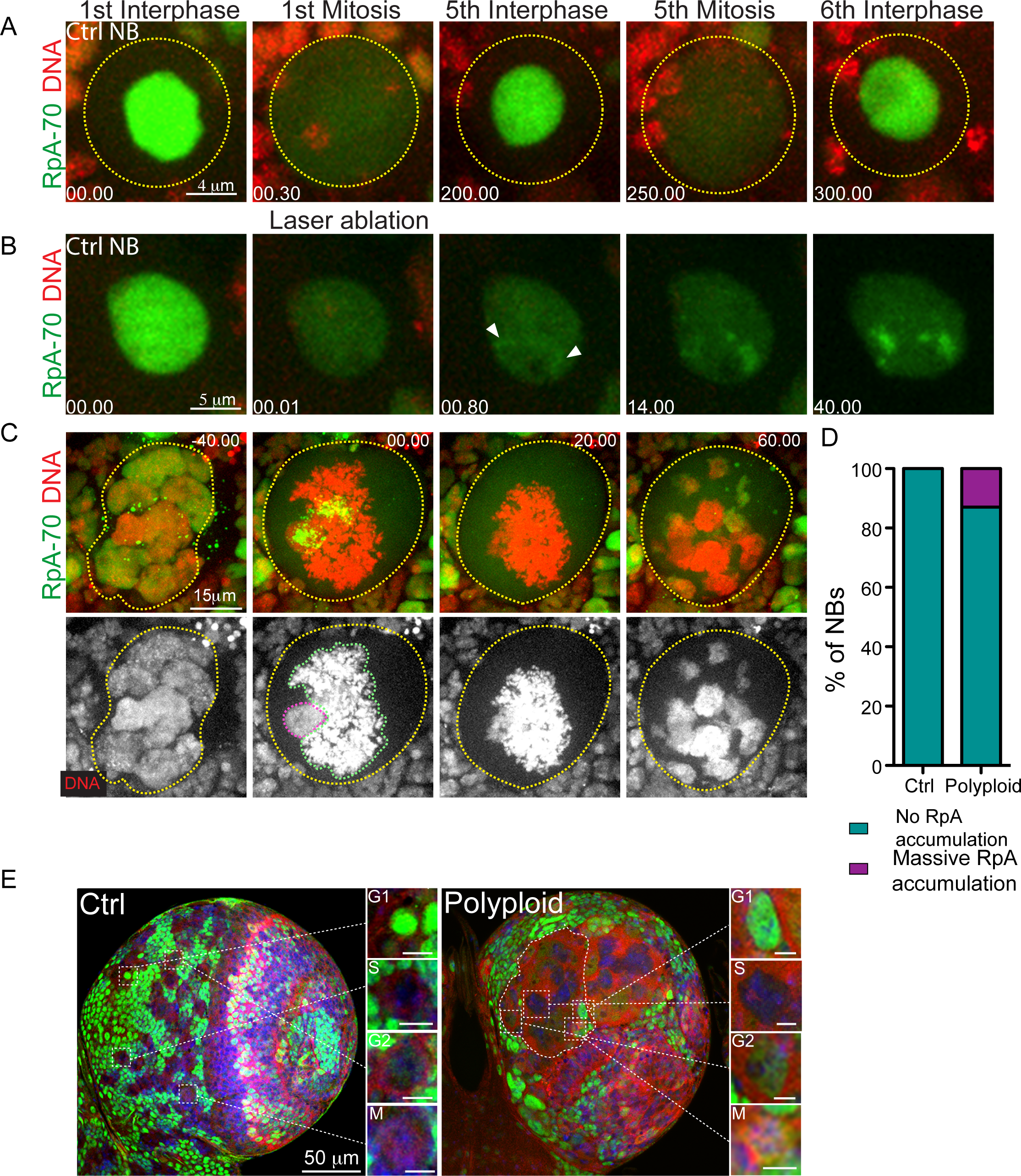
DNA damage is generated in asynchronous polyploid NBs at mitotic entry. (A-C) Stills of time-lapse movies of Ctrl (A-B) and polyploid (C) NBs expressing RpA-70-GFP to visualize ssDNA (RpA-70, shown in green) and H2Av fused to RFP to visualize DNA (in red). Yellow dashed lines outline cell boundaries, pink and green dashed lines outline nuclear boundaries. (A) RpA-70 localizes in the nucleus of interphase Ctrl NB, and becomes diffused during mitosis due to NEBD. Note that RpA-70 does not form foci during the unperturbed cell cycle of control cells, even after multiple cell cycles. (B) After laser-induced damage, RpA-70 forms foci on the damaged DNA of Ctrl NBs. (C) In this large polyploid mitotic NB a massive and transient accumulation of RpA-70-GFP on the region of less-condensed chromatin can be noticed. Time is shown in minutes.seconds. (D) Graph bar showing the percentage of Ctrl and polyploid NBs without (dark green) or with massive RpA-70-GFP accumulation at mitotic entry (violet, n=9/67 polyploid NBs). (E) Pictures of Ctrl and polyploid brain lobes expressing the FUCCI system, with GFP-E2F and RFP-CycB and stained with antibodies against GFP (in green) and RFP (in red). DNA is shown in blue. White dashed lines project magnifications of the surrounded NB, which illustrate the different cell-cycle stages. Ctrl NBs in G1-phase express only GFP in the nucleus, in S-phase express only RFP in the cytoplasm, while Ctrl NBs in G2-phase are GFP and RFP positive (mostly localized to the nucleus and to the cytoplasm, respectively). Mitotic NBs that underwent nuclear envelope breakdown exhibit diffused colocalization of GFP and RFP. Large polyploid NBs can present different nuclei at different stages of the cell-cycle, despite the presence of a common cytoplasm (n=25/105 polyploid NBs with cell cycle asynchrony).

We next analysed polyploid NBs. Surprisigly, we found that in 13.4% of polyploid NBs RpA-70-GFP massively and transiently accumulated on chromosomes upon NEBD (n=9/67 cells from 11 brains) (Figure 3C and 3D; Movie 3). Interestingly, this accumulation was restricted to few nuclei, suggesting that the generation of ssDNA was not common to all nuclei of a given polyploidy cell, consistent with the results described earlier (Figure 2B). Moreover, RpA accumulation was transient and took place during NEBD, at the prophase to prometaphase transition (Figure 3C and Movie 3).

Recently, it has been shown that replicative stress activates DNA synthesis at chromosome fragile sites (CFS) during mitosis [29]. Therefore, we investigated whether RpA recruitment in polyploid NBs was due to DNA damage occurring at mitotic entry or was identifying sites of DNA synthesis. To analyse DNA replication, we incubated NBs with 5-Ethynyl-2’-deoxyuridine (EdU), a thymidine analog which is incorporated into DNA during DNA replication, and used Ph3 to identify mitotic cells (Supplementary Figure 3A). We limited EdU incorporation to 1h, since NBs have extremely short cell cycles (1.6h on average elapsed from one mitosis to the next [30] and our work). Edu levels in interphase (Ph3-) were higher in Ctrl than in polyploidy NBs (Ctrl: 1482±136.3, n=30 NBs and polyploid: 901.3±111.9, n=30 NBs, p=0.0017) (Supplementary Figure 3B-C). Importantly, however, the levels of EdU in mitosis were much lower (Ctrl:239.3±82.5, n=30 and polyploid: 183.9±10.6) and we only detected one EdU positive mitotic (Ph3+) Ctrl NB, which most likely reflects a short G2 phase (Supplementary Fig. 3B and D). We then analysed Ph3+ polyploid NBs that contained γ–H2Av foci, indicative of damage, reasoning that - if DNA synthesis was occurring - these nuclei should be EdU positive. However, this was not the case as we never detected EdU in mitotic, γ–H2Av positive polyploid NBs (Supplementary Figure 3D). We concluded that in NBs, DNA synthesis does not occur during mitosis and the ssDNA foci that appear in a small portion of polyploid NBs during mitotic entry most likely reflect the generation of DNA damage at this cell cycle stage.

### Cell cycle asynchrony triggers DNA damage in polyploid mitosis

While filming NBs expressing RpA, we noticed that the nuclei that accumulated RpA foci were delayed in mitotic entry when compared to neighbouring nuclei of the same polyploid NB. Indeed, their DNA appeared less condensed than the DNA of neighbouring nuclei (Figure 3C and Movie 3). Importantly, asynchronous mitotic entry was also observed in polyploid NBs induced through Pav KD as described in [22].

The lack of synchrony at mitotic entry, suggests that cell cycle progression is not synchronous in all nuclei of polyploid cells. To investigate this possibility, we used the Fly-FUCCI system [31]. The Fly-FUCCI allows to discriminate between the different phases of the cell cycle by expressing truncated forms of E2F and Cyclin B (CycB) fused to GFP and RFP respectively (GFP-E2F 1-230, RFP-CycB 1-266). We found that 23.8% of polyploid NBs were characterized by the presence of a mixed population of nuclei at different cell cycle stages (n=25/105 NBs from 10 brains; Figure 3E). It is important to mention that we could also detect polyploid NBs in which certain nuclei of a given cell entered and exited mitosis while other nuclei of the same cell remained in interphase (Supplementary Figure 4A), as assessed by the absence of NEBD. This type of behaviour, which we named complete asynchrony, was observed in 12.7% of polyploid NBs (16/126 NBs from 12 brains) (Supplementary Figure 4A-B).

**Figure 4.**
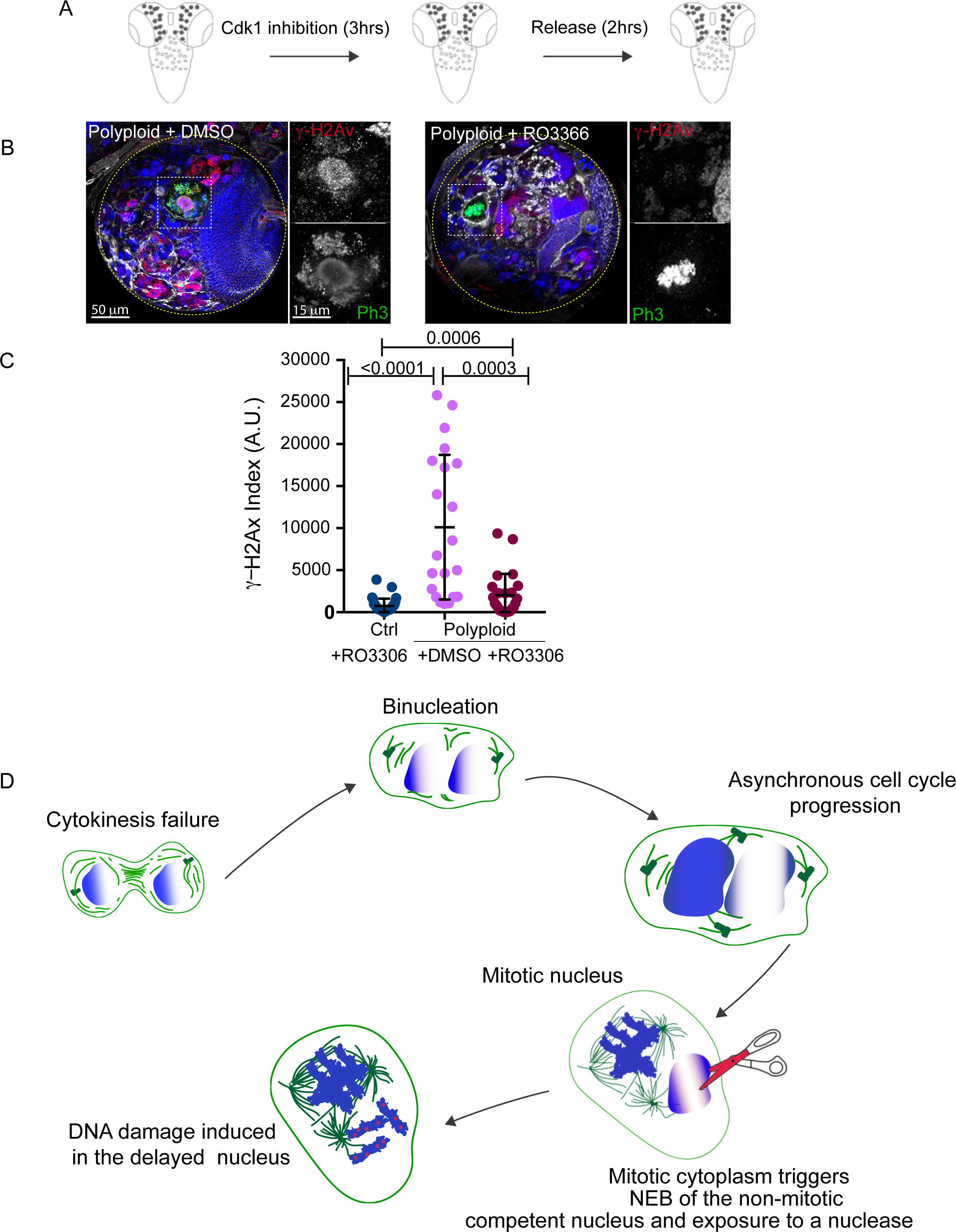
Forcing cell cycle synchronization attenuates DNA damage in mitotic NBs. (A) Schematic representation of the experimental strategy used to enforce cell cycle synchrony in *Drosophila* larval brains. Brains were incubated for 3 hrs with the Cdk1 inhibitor RO3306 and released from the inhibition for 2 hrs. (B) Micrographs of polyploid brains treated with DMSO (left) or RO3306 (right) and stained with antibodies detecting DNA double-strand breaks (γ-H2Av, in red in the merged panel and in gray) and Ph3 (which labels mitotic cells, in green in the merged panel and in gray). The yellow dashed lines outlines the brain lobes. White dashed squares surround the NBs magnified in the insets. Note the lack of DNA damage in mitotic NBs that were treated with RO3306. (C) Dot plot charts showing the γ-H2Av index in mitotic Ctrl NBs treated with RO3366 (blue), polyploid NBs treated with DMSO (magenta) and polyploid NBs treated with RO3306 (dark red). Statistical significance was assessed by unpaired t-test. (D) Schematic diagram of the model proposed. For simplicity, polyploid cells were drawn as being binucleated. After cytokinesis failure, polyploid cells lose synchrony in cell cycle progression. The nucleus that is competent to undergo mitosis (left) will undergo NEBD, which will trigger mitosis entry in the yet “unready” nuclei. This second nucleus (right) is exposed to a mitotic cytoplasm that might contain nucleases that can act on non-condensed chromatin, generating DNA damage.

The restriction of DNA damage to a single nucleus (Figure 2H) of binucleated mNSCs also implied cell cycle asynchrony in these cells. Since mNSCs cells did not survive FUCCI transfection, we decided to characterize cell cycle stages using Proliferating Cell Nuclear Antigen (PCNA), which distinctively labels S-phase and Ph3. In NT or EtOH treated cells, we observed individual nuclei positive for either PCNA (in S-phase), Ph3 (mitotic) or negative for both (Supplementary Figure 4C). In LatB treated cells, however, 11.0% of binucleated cells were asynchronous (n=35 out of 320), as evidenced by the presence in the same cell of one nuclei positive for PCNA (end of S phase) and one nuclei negative for PCNA, but Ph3 positive (Supplementary Figure 4D-E). These results suggest that cell cycle asynchrony is a characteristic of cells with more than one nucleus and increased DNA content.

We next tested whether cell cycle asynchrony was indeed other to DNA damage at mitotic entry. After several genetic and chemical tests (not shown), we choose to use the Cdk1 RO3306 inhibitor, which supresses mitotic entry. We assessed the efficacy of RO3306 treatment by scoring for Ph3 positive cells, which were absent from both Ctrl and polyploid brains upon RO3306 treatment (not shown). However, after washout followed by 2hrs release, mitotic cells were readily identified (Figure 4A). Strikingly, the levels of DNA damage in mitotic cells were considerably decreased in polyploid NBs, when compared with polyploid cells incubated with DMSO (Polyploid+DMSO 10114±1877, n=21 NBs, Polyploid+RO3306: 2812±612.2, n=24 NBs, p=0.0003) (Figure 4B-C). Condensed metaphase-like chromosomes lacking γ-H2Av signals were noticed, suggesting that these cells have entered mitosis without generating any damage.

These findings suggest that blocking cell cycle progression allows synchronization between different nuclei of polyploid NBs. Further, synchronization prevents forced mitotic entry of yet not competent nuclei and decreases the likelihood of generating DNA damage at mitotic entry.

In this work, we have analysed the consequences of polyploid cell proliferation and we have identified a novel way of generating genetic instability at mitotic entry. Our findings are compatible with a model where a fraction of polyploid (multinucleated or binucleated) cells progress asynchronously through the cell cycle (Figure 4D). We report the co-existence of mitotic and non-mitotic nuclei in polyploid cells. We show that mitotic nuclei can force neighbouring, not yet mitotic competent-nuclei to undergo NEBD and rapid transition through mitosis. This leads to chromosome segregation errors as shown in [22], but also to DNA damage generation during mitosis (this work). By which means the mitotic cytoplasm of neighboring nuclei can trigger DNA damage in delayed nuclei, remains to be explained. A likely possibility is the presence of nucleases that can act on uncondensed chromatin. Although future work will be required to address this question and identify putative DNA nucleases, enzymes such as TREX1 [32] or LEM-3 [33] appear as good candidates. These nucleases are active and present in mitotic cytoplasms and play essential roles in resolving chromatin bridges. It is tempting to speculate that they could recognise poorly condensed, non-mitotic nuclei in polyploid cells.

It is important to mention that this way of generating DNA damage and chromosome instability can not solely account for the high levels of DNA damage observed in polyploid NBs. Indeed, only a small fraction of polyploid NBs and mNSCs present DNA damage in mitosis restricted to a single nucleus. Moreover we found that forced mitotic entry of unready nuclei, can only occur in a small fraction of polyploid cells, even if cell cycle asynchrony is present. These results suggest that even if rare, such an event might contribute to severe genome reshuffling found after WGDs [2–6]. It will be interesting to determine what type of chromosome rearrangements can be identified after such catastrophic events.

Moreover, it remains to be investigated what is the role of transisent RpA accumulation upon DNA damage generation at mitotic entry, and whether this type of damage can be repaired in the following cell cycle.

Classic cell fusion experiments [34–36] have shown that the mitotic cytoplasm is normally dominant and triggers premature chromosome condensation in the neighbouring interphase nuclei. The work presented here sustains this view and show that forced mitotic entry might represent a source of DNA damage when chromosomes are not condensed. Overall, this work and the study from [22] provide insight into how genetic instability can be established in polyploid cells. They shed light on a complex plethora of events underlying the generation of DNA damage in polyploid cells. Our findings will help laying the foundation of new studies, which will establish the contribution of whole genome duplications to genetic instability and cancer.

## Supporting information

Nano et al Sup info

## Acknowledgements

We thank D. Gogendeau, S. Gemble, M. Rujano, D. Gambarotto, D. Vargas, D. Fachinetti, A.J. Bardin. A. Echard, A. Goupil and F. Edwards for helpful discussions and critical reading of the manuscript. We thank A.J. Bardin and Y. Bellaiche (Institut Curie, FR), R. Karess and A. Guichet (IJM, FR), C. Lehner (UZH, CH), J.M. Abrams (UT Southwestern, TX), E.F. Wieshaus. (Princeton University, NJ), C.Q. Doe (HHMI, OR) and J. Raff (Oxford University, UK) for sharing flies and reagents. We thank V. Racine from QuantaCell for helpful discussions on data analysis and L. Sengmanivong and L. Leconte for help with confocal microscopy. We acknowledge the Nikon Imaging Centre at Institut Curie-CNRS, member of the French National Research Infrastructure France-BioImaging (ANR10-INSB-04) for equipment and support, the Bloomington Drosophila Stock Centre (NIH P40OD018537) at Indiana University, USA, the Developmental Studies Hybridoma Bank at University of Iowa (DSHB, IA, USA) and the Therapeutic Antibodies and Recombinant Antibodies Platform at Institut Curie (FR) for DNA clones, *Drosophila* stocks and reagents. This work was supported by an ERC CoG (ChromoNumber - LS3, ERC-2016-COG), the LNCC, Comite de Paris (24097-2017), Institut Curie, INSERM for V.M. and the CNRS for R.B and the Basto team. M.N. was supported by grants from the Ligue nationale contre le cancer and FRM (FDT20160435352). Our lab is a member of the Labex CelTisPhyBio.

## Author contribution

M.N. and R.B conceived the project, analysed the data and wrote the manuscript. M.N. did most of the experimental procedures. C.P. generated tools. V.F. helped with laser ablation experiments. A.S. and V. M. performed the experiments and quantifications in mNSCs. R.B. supervised the project.

