## Supplementary material for "Cell Cycle Asynchrony Generates DNA Damage at Mitotic Entry in Polyploid Cells": Nano et al Sup info

### Supplementary Figure 1- Nano et al.

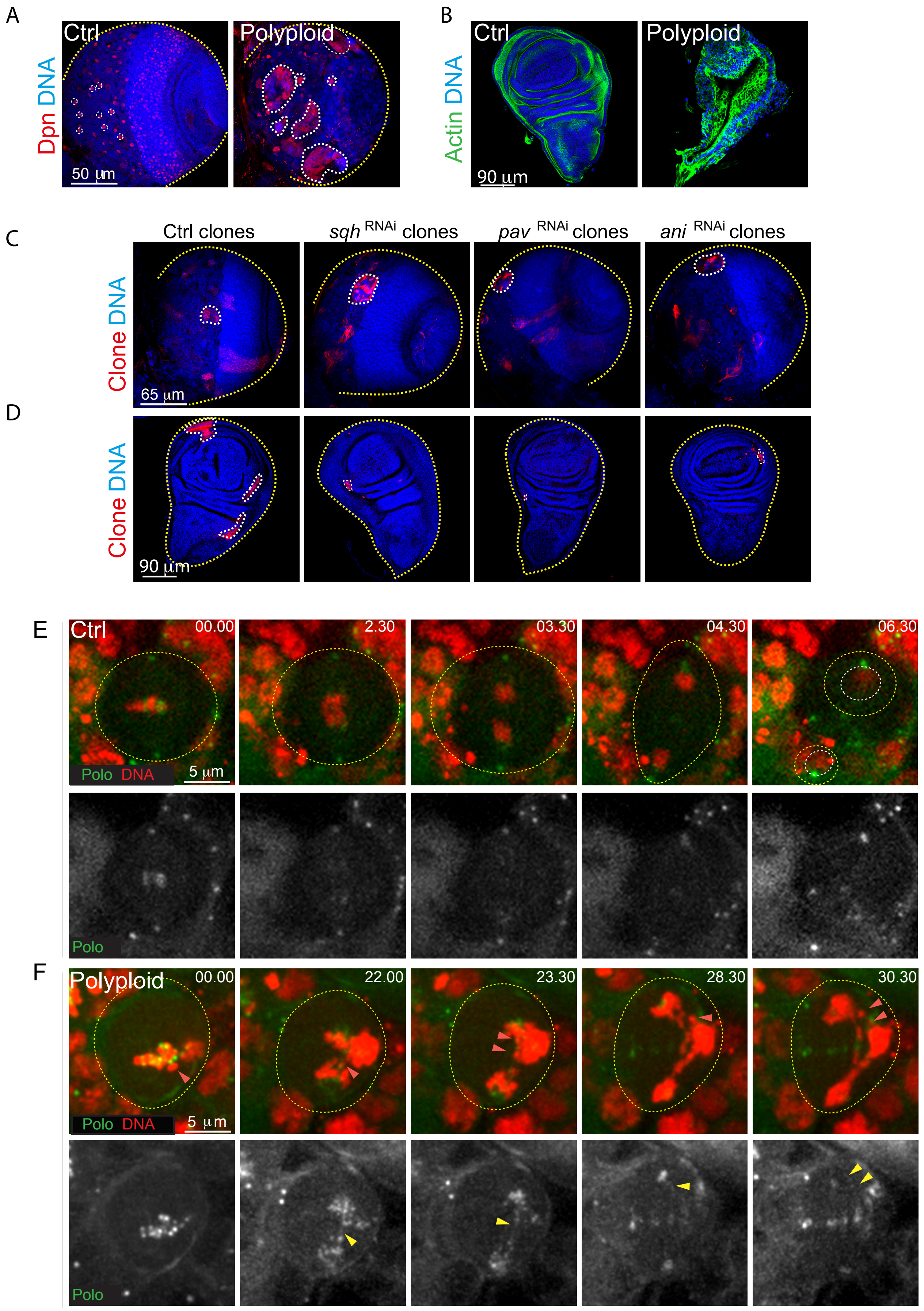

### Supplementary Figure 2- NANO et al

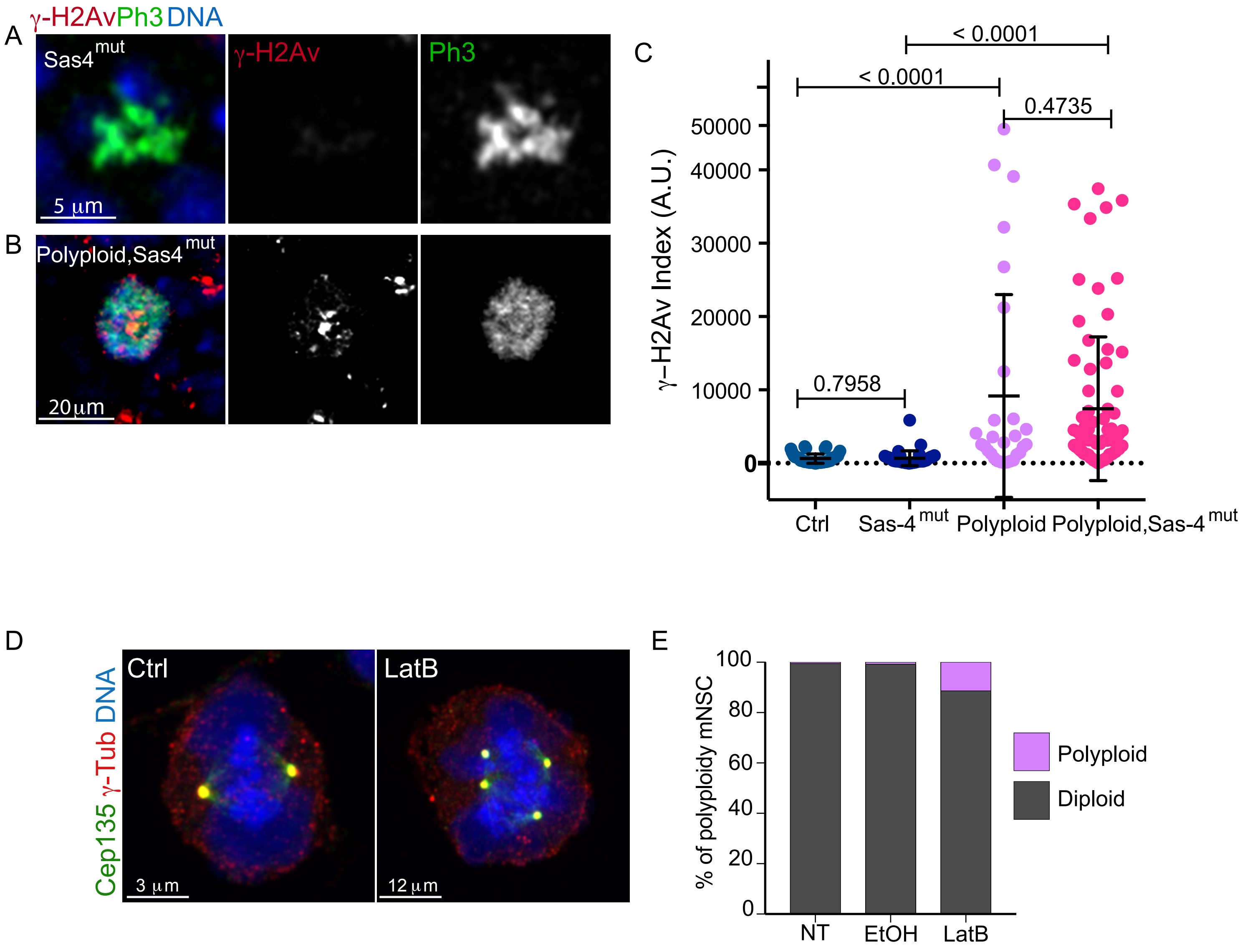

### Supplementary Figure 3- NANO et al

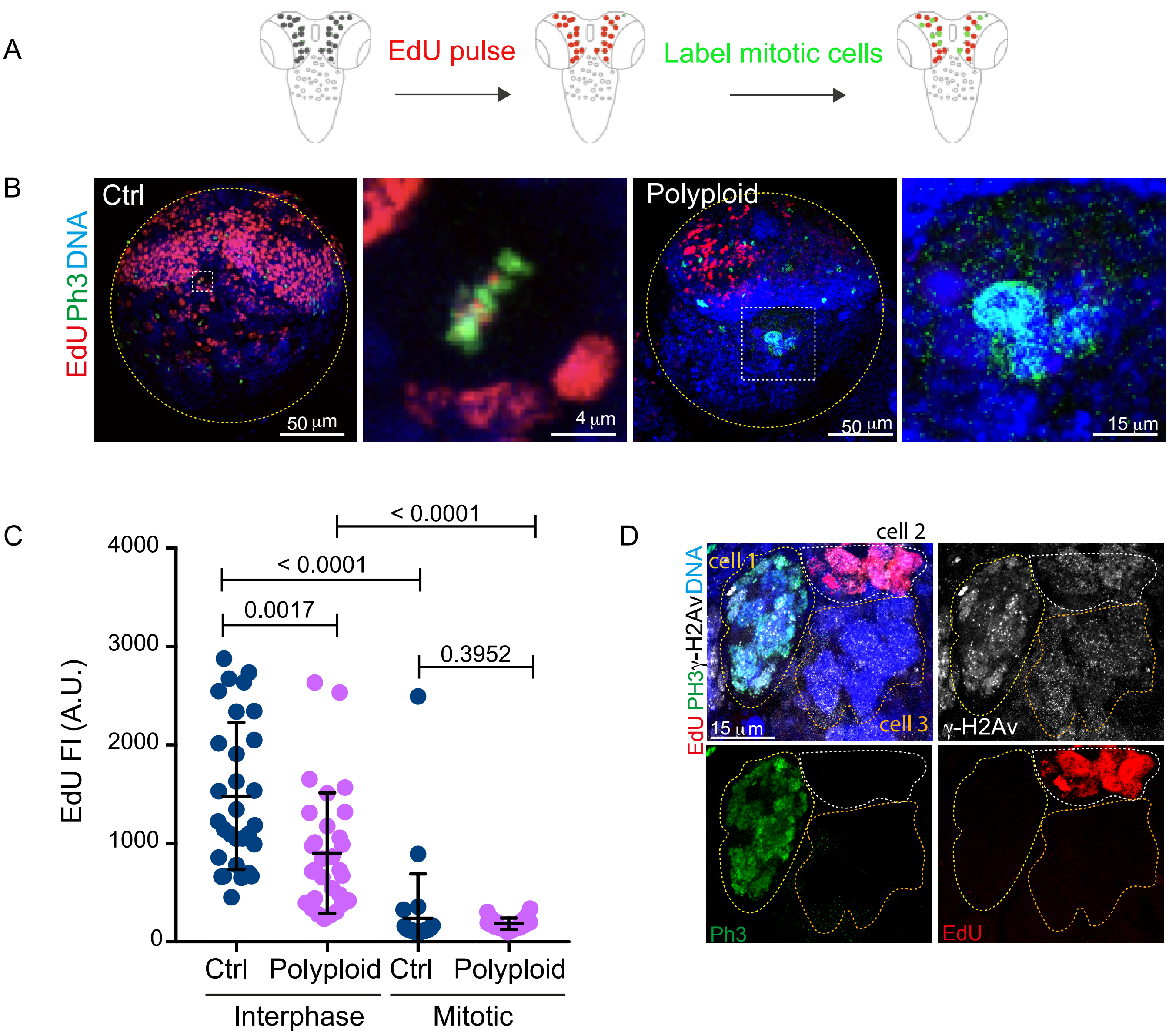

### Supplementary Figure 4- NANO et al

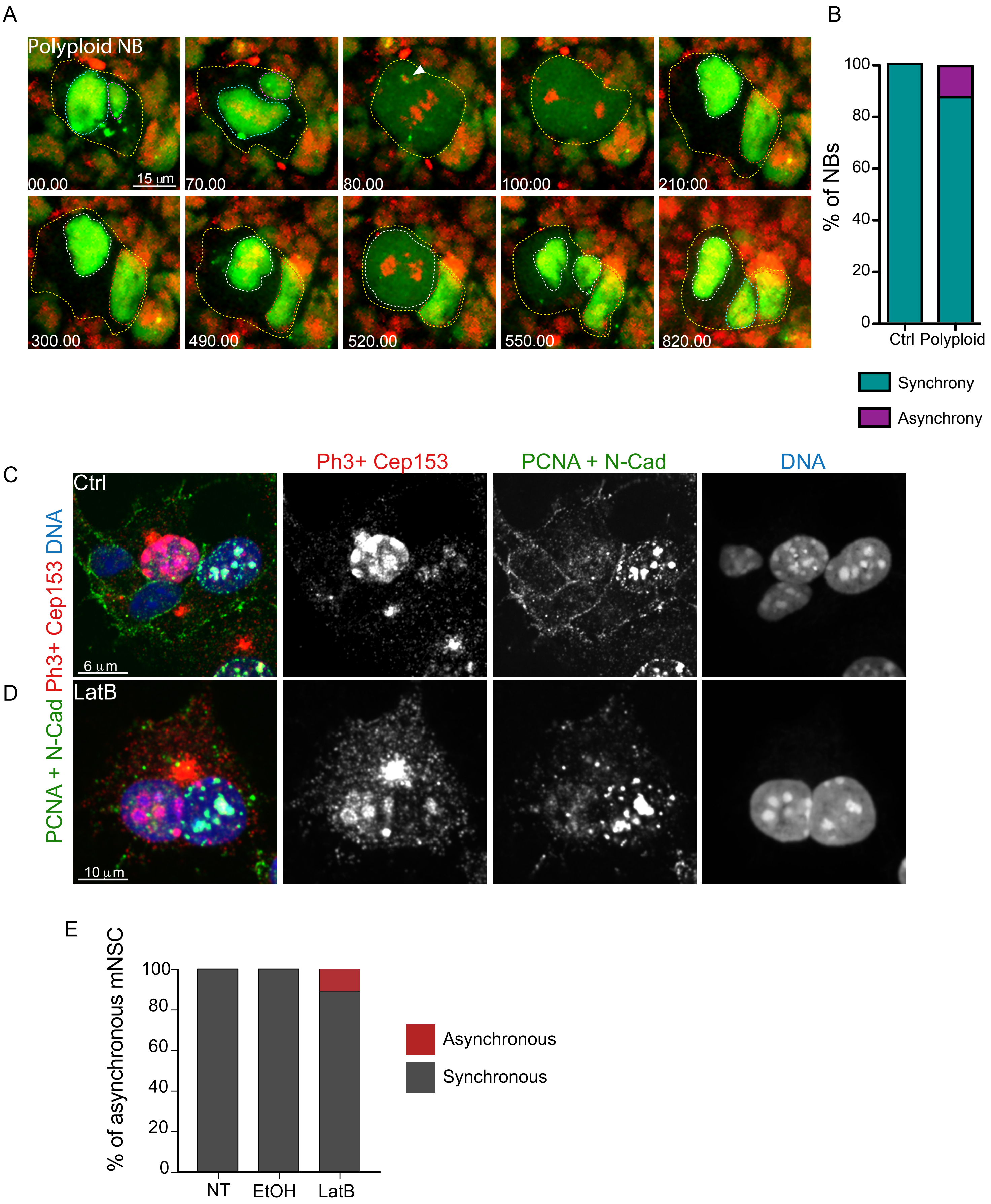

### Supplementary Information

### CELL CYCLE ASYNCHRONY GENERATES DNA DAMAGE AT MITOTIC ENTRY IN POLYPLOID CELLS

#### Supplementary Figure legends

**Supplementary Figure 1. Tissues that tolerate polyploidy in developing *Drosophila* larvae remain proliferative despite the presence of structural chromosomal aberrations.** (A) Representative picturs of Ctrl and polyploid brain lobes labeled with Dpn antibodies (red) to label NBs, showing that polyploid NBs have different sizes and maintain neural stem cell fate. Yellow dashed lines label brain lobes and white dashed lines label individual diploid (in Ctrls) or polyploid NBs. (B) Representative pictures of Ctrl and polyploid wing discs, labeled with phalloidin (green) showing that polyploid wing disc do not grow as Ctrl wing discs. (C-D) Representative pictures of Ctrl, $sqh^{RNAi}$, $pav^{RNAi}$ and $ani^{RNAi}$ brain clones (top) and wing disc clones (bottom). The clone represents a group of cells where a genetic alteration has been induced. It is labeled with a membrane-localized RFP and stained with antibodies against RFP. DNA is shown in blue. Non-RFP cells are wild type cells. Yellow dashed lines outline brain lobe and wing disc boundaries and white dashed lines highlight the clones. Note that, in the brain, polyploid clones are much larger size than the clones in the wing disc. (E-F) Stills of time-lapse movies of mitotic NBs expressing H2Av fused to RFP to visualize DNA (shown in red in the merged panels) and Polo fused to GFP (shown in green in the merged panels and in grey). Yellow dashed lines outline cells boundaries. Time is shown in minutes.seconds. (E) Representative mitosis of a Ctrl NB. Polo localizes on centrosomes, kinetochores, and at the level of the ingressing cytokinetic furrow. White dashed lines outline the daughter nuclei formed at the end of mitosis. (F) Polyploid NB mitosis. Polo can be seen localizing on centrosomes and kinetochores. In addition, fine threads of Polo (yellow arrowheads) can be seen linking acentric chromosomes (red arrowheads) to the main chromosome mass (n=11/11 NBs).

**Supplementary Figure 2. Multinucleation does not greatly contribute to DNA damage generation during mitosis in polyploid NBs**

(A-B) Representative micrographs of $Sas4^{mut}$ and polyploid, $Sas4^{mut}$ NBs labeled with antibodies against γ-H2Av (red) and Ph3 (green). Mitotic $Sas4^{mut}$ NBs do not show DNA damage, while polyploid, $Sas4^{mut}$ NBs with a single nucleus can still show DNA damage in mitosis. (C) Dot plot chart of γ-H2Av index in Ctrl (blue), $Sas4^{mut}$ (dark blue), polyploid (magenta) and polyploid, $Sas4^{mut}$ NBs (pink). (D) Micrograph of Ctrl and LatB-treated mNSCs. mNSCs were labelled with Cep135 (in green) and γ-Tubulin (in red) to unambiguously identify centrosomes. DNA is shown in blue. Note how LatB mNSC accumulate multiple centrosomes due to cytokinesis failure. (E) Graph bar showing the frequency of diploids (in gray) and polyploid cells (magenta) in the mNSC population. Statistical significance was assessed by unpaired t-tests.

**Supplementary Figure 3. Analysis of EdU incorporation in mitotic polyploid NBs and DNA damage in polyploid NBs without centrosomes (**A) Schematic representation of the experimental strategy used to monitor DNA synthesis during mitosis in *Drosophila* larval brains. Brains were incubated with EdU for 1 hr *ex-vivo* and were then labelled with Ph3 to identify mitotic cells. (B) Micrograph of Ctrl and polyploid brain lobes and NBs labelled with EdU (in red) and with antibodies directed against the mitotic marker Ph3 (in green). Yellow dashed lines outline the lobe boundaries, white dashed lines highlights the NBs magnified in the inset. Note that we only scored one mitotic Ctrl NB positive for EdU, suggesting that the incorporation of EdU did not take place during mitosis. (C) Dot plot chart showing mean EdU fluorescence intensity (FI, expressed in arbitrary units – A.U.) measured in interphase and mitotic Ctrl NBs (blue) and polyploid NBs (magenta). (D) Micrograph of polyploid NBs labelled with EdU (in red), with antibodies directed against the mitotic marker Ph3 (in green) and with antibodies detecting DNA double strand breaks (γ-H2Av, in gray). Note how DNA damage in the mitotic NB (cell 1) is not associated with EdU incorporation, indicating that the presence of DNA damage markers in mitotic cells reflects bona-fide DNA damage rather than products of DNA synthesis.

**Supplementary Figure 4. Asynchronous cell cycle progression in polyploid NBs and mNSCs.**

(A) Stills of time-lapse movie of polyploid NB expressing RpA-70-GFP to visualize ssDNA (green) and H2Av fused to RFP to visualize DNA (in red), undergoing multiple cell cycles. An initially binucleated NB enters mitosis at t=80.00. Note how RpA-70 forms foci on the damaged DNA. At t=210.00, the polyploid NB exits mitosis and forms two new nuclei. At the following mitotic entry (t=520.00), only one of the nuclei undergoes NEBD, while the other remains in interphase (the nuclei are outlined by a white and a red dotted line, respectively). We named this phenomenon complete asynchrony, to distinguish it from the case where delayed nuclei are ultimately forced to enter mitosis. Time is shown in minutes.seconds. (B) Graph bar showing the percentage of Ctrl and polyploid NBs experiencing cell cycle synchrony (dark green) and complete asynchrony (violet). (C-D) Micrographs of Ctrl and LatB mNSCs labelled with antibodies detecting PCNA to identify cells in S-phase and N-Cadherin (N-Cad) to label cell membrane (in green) and Ph3 and Cep153 (in red). DNA is shown in blue. (E) Graph bar showing the frequency of synchronized (in gray) and asynchronous (in red) cells in the mNSCs population.

**Methods**

CONTACT FOR REAGENT AND RESOURCE SHARING

**Animals**

**Fly husbandry and fly stocks**

Flies were raised on cornmeal medium (0.75% agar, 3.5% organic wheat flour, 5.0% yeast, 5.5% sugar, 2.5% nipagin, 1.0% penicillin-streptomycin and 0.4% propionic acid). Flies stocks were maintained at 18°C. Crosses were carried out in plastic vials and maintained at 25°C unless differently specified. Stocks were maintained using balancer inverted chromosomes to prevent recombination. Stocks used in this study: $sqh^1$ [1], H2Av-mRFP (BL#23651) [2], Spc-25-GFP [3], Polo TRAP [4], RpA-70-GFP [5], FUCCI#2 (BL#5509) [6], $sqh^{RNAi}$ (BL#32439) [7], $anillin^{RNAi}$ (BL#53358) [8] and $pavarotti^{RNAi}$ (BL#42573) [9]. To induce clones, we used as a driver *hsflp*;;$Tub^{FRT}GAL80^{FRT}$GAL4,UAS-mCD8-RFP [10]. BL=Bloomington *Drosophila* Stock Center (Indiana University, IN, USA). The $sqh^1$ recessive allele is located on the X chromosome and in this study it was balanced with an FM7 chromosome or with an FM7 chromosome that contained the Kruppel-GAL4, UAS-GFP marker (FM7, Kr)[11] to facilitate the selection of hemizygous mutant larvae (always males). Heterozygous female larvae or hemizygous FM7 or FM, Kr males were used as diploid controls (Ctrl). Further, when using drivers, Ctrls also included the chromosomes containing the drivers. We also used other Ctrl stocks such as: *wf* and WT isogenic for $sqh^1$ mutant (WT iso: *wf* crossed with males FM7, Kr and propagated for several generations).

**Fly staging**

In most experiments except FISH (Figure 1E), and Deadpan staining (SFigure 1A), larvae were staged to obtain comparable stages of development. Egg collection was performed at 25°C for 24 hours. Clone induction was carried out by heat shocks of $1^{st}$ instar larvae for 45min at 37°C. After development at 25°C, third instar larvae were used for dissection. To compensate for developmental delay, $sqh^1$ brains one day older were dissected concomitantly to Ctrl brains.

**Brain overgrowth**

Brain overgrowth was assessed comparing the latest stage of third-instar Ctrl WTiso larvae (dissected at day 6 of development) and *sqh*$^{1}$ aged larvae (dissected at day 13 of development).

**Mice.** For animal care the European and French National Regulation for the Protection of Vertebrate Animals was followed for experimental and other Scientific Purposes (Directive 2010/63; French Decree 2013-118). The project was authorized and benefited from guidance of the Animal Welfare Body, Research Centre, Institut Curie. C57Bl6/N pregnant females were purchased from Charles River Laboratories (France). For all pregnant females, the day of mating was referred to 0.5 day of gestation (E0.5). Pregnant females were anaesthetized by inhalation of isoflurane (Baxter, DDG9621) before being sacrificed by cervical dislocation. Embryos were collected after caesarian section.

**Primary adherent MNSC culture and drug treatment**

To establish primary adherent cultures of embryonic mNSCs we have adapted the protocol described in [12]. E16.5 cerebral cortex explants were dissociated into a single cell suspension after 5 min accutase (Sigma, A6964) incubation at 37°C. Cortical cells were plated at a density of $1x10^4$ cells/cm2 on plastic pre-coated sequentially for 1h at RT with poly-D-lysine (2µg/cm$^2$) and fibronectin (1µg/cm$^2$). Cells were cultured in a medium favoring the propagation of proliferating cells (proliferation medium) containing DMEM/HamF12 medium, supplemented with glucose (2.9g/L), sodium bicarbonate (1.2g/L), B27 supplement w/o vitamin A (2%), FGF (10 ng/ml), EGF (10 ng/ml), penicillin/streptomycin (1%) and Amphotericin B (250ng/ml), at 37°C under 5% $CO_2$. After one week in culture, the population was only composed of nestin-positive proliferating NSCs (data not shown).

Embryonic NSCs were seeded at $1x10^4$ cells/cm$^2$ on glass coverslips pre-coated with poly-D-lysine and fibronectin in proliferating medium. The day after, cells were treated for 30 min at 37°C with ethanol (drug diluent, 0.015%) or latrunculin B (5µM, Millipore, 428020) before 2 quick washes in proliferation medium. The efficiency of drug treatment was assessed by cell rounding and was considered as time 0 (T0). Cells were enabled to proliferate in proliferation medium for 16h before PFA or methanol fixation and immunofluorescent staining. For EdU incorporation experiments, cells were incubated with

EdU for 5 min before PFA fixation according to Manufacturer' instructions (Click-IT EdU Alexa Fluor 647 Imaging Kit, Molecular Probes, C10640).

**Whole mount tissue preparation and imaging of *Drosophila* larval brains**

Brains from third instar larvae were dissected in PBS and fixed for 25 minutes in 4% paraformaldehyde in PBS. They were washed three times in PBST 0.3% (PBS, 0.3% Triton X-100 (T9284, Sigma), 10 minutes for each wash) and incubated several hours in agitation at room temperature (RT) and O/N at 4 °C with primary antibodies at the appropriate dilution in PBST 0.3%. Primary antibodies used in this study are: rabbit anti-γ-H2Av (1:500; Rockland#600-401-914, Limerick, PA, USA), rabbit anti-Ph3 (1:250; Sigma-Aldrich#H6409, Merck, Darmstadt, DE), guinea pig anti-Deadpan (generated in our lab [13]), human anti-GFP (1:250; Antibody platform IC-A-R-H#11, Paris, FR), rabbit anti-GFP (1:250; TermoFisherScientific#A-11122, Waltham, MA, USA), mouse anti-GFP clone JL-8 (1:500; Clontech#632381, Mountain View, CA, USA), chicken anti-GFP (1:500; Abcam# ab13970; Cambridge, UK), human anti-m-Cherry (1:100; Antibody platform IC-A-R-H#46) and rat anti-RFP clone 5F8 (1:500; ChromoTek, Martinsried, DE). Tissues were washed three times in PBST 0.3% (10 minutes for each wash) and incubated several hours in agitation at RT and O/N at 4 °C with 3 μg/ml DAPI to label DNA (4',6-diamidino-2-phenylindole; 62248, Termo Fisher Scientific) and secondary antibodies diluted in PBST 0.3%. All secondary antibodies (conjugated Alexa Fluor 488, Alexa Fluor 546 and Alexa Fluor 647) and Phalloidin 488 to label actin were used at 1:250 and were purchased from Molecular Probes (Invitrogen, TermoFisherScientific). Brains were then washed three times in PBST 0.3% (10 minutes for each wash), rinsed in PBS and mounted in mounting media. Standard mounting media was prepared with 1.25% n-propyl gallate (Sigma, P3130), 75% glycerol (bidistilled, 99.5%, VWR, 24388-295), 23.75% $H_2O$.

Images were acquired on a confocal Nikon A1R inverted Ti-E microscope with oil 40X NA 1.3 or 60X NA 1.4 objectives in NIS Element software. Interval for z-stacks acquisitions was set up from 0.3 μm to 1 μm. Images were analysed with Fiji. The cellular structures of interest were quantified by manual scoring of the individual slices for each z-stack. If quantitative analysis of the fluorescent signal was required, settings were kept constant for all the acquisitions. Statistical analysis was performed with Graph Pad Prism using the tests mentioned in the figure legends. Further details on how DNA damage was quantified are available below.

**Fluorescence In Situ Hybridization (FISH)**

FISH was used to assess the copies of chromosome II (Chr II) in Ctrl cells or cells that fail cytokinesis and was carried out as described in [13]. Briefly, oligonucleotide probes for AACAC repeats (chromosome II) [14] were synthesized with a 5'CY3 fluorescent dye (sequence: 5' CY3-AACACAACACAACACAACACAACACAACACAACAC) (GenScript, George Town, KY). 3rd instar larval brains were dissected in PBS and fixed 30 minutes in 4% paraformaldehyde in PBS 0.1% Tween-20 (162312, Panreac Sintesis, Barcelona, ES). Tissues were washed three times in PBS, once in 2XSSC (EU0300, Euromedex, Souffelweyersheim, FR)/0.1% tween-20 (2XSSCT) and once in 2XSSCT/50% formamide (47671, Sigma, St. Louis, MO, USA). Brains or WDs were transferred in a PCR tube containing 2XSSCT/50% formamide at 92°C for pre-hybridization and denaturated 3 minutes. Tissues were then hybridized 5 minutes at 92°C with the previously denaturated DNA probe (40 ng) in hybridization buffer, composed of 20% dextran sulphate (D8906, Sigma), 2XSSCT/50% deionized formamide (F9037, Sigma) and 0.5 mg/ml salmon sperm DNA (D1626, Sigma). Hybridization was allowed to proceed O/N at 37°C. Brains or WDs were then washed once with 2XSSCT at 60°C for 10 minutes and once for 5 minutes at room temperature. Samples were rinsed in PBS and labelled with DAPI (in PBS 0.1% Triton X-100) for at least 30 min. Tissues was mounted using standard mounting medium. Image acquisition was made as described for whole mount preparation of tissues and FISH signals were quantified manually for each z-stack.

**EdU incorporation assay**

To analyse S-phase in polyploid NBs, brains were incubated with EdU. 3rd instar larval brains were dissected in Schneider's *Drosophila* Medium (described in the Methods section relative to live imaging) and incubated for 1 hour at 25°C in the same medium with 100 μM EdU (5-ethynyl-2'-deoxyuridine). Brains were then washed in PBS, fixed and immunostained. EdU detection was performed after secondary antibody detection, according to the manufacturer instructions (C10640, Molecular Probes, TermoFisherScientific, Waltham, MA, USA). Image acquisition and treatment were made as described for whole mount preparation of tissues. Quantitative analysis of EdU nuclear coverage was performed as described below.

**RO3306 treatment**

*Drosophila* larval brains were dissected and incubated 3hrs *ex-vivo* in Schneider's *Drosophila* Medium containing either 500 μM of the Cdk-1 inhibitor RO3306 or DMSO as a control. Brains were washed 3X and incubated in fresh Schneider's *Drosophila* Medium for 2 hrs before being rinsed in PBS, fixed and immunostained as described for whole mount tissue. Image acquisition and treatment were made as described for whole mount preparation of tissues. Quantitative analysis of γ-H2AV fluorescemce intensity and nuclear coverage was performed as described below.

**Quantitative analysis of DNA damage and EdU coverage in single cells – *Drosophila* NBs**

Staged 3$^{rd}$ instar larval brains were dissected, stained and imaged using the procedures described above. We used the γ-H2Av primary antibody, which was preferentially detected using a secondary antibody conjugated Alexa Fluor 546, which was found to give the best signal to noise ratio. For evaluation of EdU nuclear coverage, EdU was detected with Alexa Fluor 647. Imaging was performed using a 40X NA 1.3 oil objectives in NIS Element software with z-stacks of 1μm. To evaluate fluorescence intensity, excitation parameters for the channel of interest were maintained constant. Image analysis was performed using Fiji. Images were imported as z-stack of 3 channels, being the red channel the one subject to analysis. Nuclei were manually segmented using the *Freehand selection* tool in their central z plane (determined on DAPI signal) and each nucleus was saved as a region of interest (ROI). During the segmentation process, the channel of interest was not taken into account, to avoid any possible bias. Micronuclei were not analysed. Quantitative values were obtained using a new plugin developed by QUANTACELL (www.quantacell.com). The channel of interest was segmented to separate positive pixels from negative pixels by a thresholding operation. After a number of tests, we assigned a constant threshold value (562 for measurement of DNA damage, 568 for EdU analysis). The minimum size of objects was set to one pixel. The plugin generates an output for the segmented ROIs indicating, for each nucleus: z position, area in μm$^2$, the percentage of positive pixels (named *coverage*) and the average intensity of the channel of interest for the ROI (named *fluorescence intensity*-FI). Additionally, the plugin produces a *montage*, which is an RGB image combining all cells analysed in the z-stack, providing an overview of the measured ROIs. Once obtained the output, all values of coverage and FI were compared visually with the montage, to ensure consistency.

**Immunofluorescent and image analysis of mNSCs**

Cells were fixed for 20 min in PFA 4% at RT or for 8 min at -20°C in methanol. Cells were permeabilized for 10 min in PBS, TX100 0.5%, incubated for 30 min in blocking solution (PBS, BSA 2%, Triton X-100 0.1%, Sodium azide 0.02%) before sequential incubation for 2h at RT with primary and secondary antibodies diluted in the blocking solution. Primary antibodies used are listed in Table 1. All secondary antibodies (highly-crossed absorbed conjugated Alexa Fluor 488, Alexa Fluor 546 and Alexa Fluor 647) were used at 1/500 dilution and purchased from Molecular Probes (Invitrogen). 4',6-diamidino-2-phenylindole (DAPI, ThermoFisher Scientific) was used to label DNA at a concentration of 3mg/ml. Images were acquired on a Leica epifluorescence microscope equipped with a sCMOS Hamamatsu camera with a 100X NA1.40 objective or on a Gattaca/Nikon wide-field spinning disk confocal microscope equipped with a sCMOS Prime95B Photometrics camera with a 100X NA 1.40 objective.

The incidence of polyploidy in the different conditions was assessed by evaluating the number of centrosomes in mitotic NSCs at the prometaphase stage based on a co-staining with two centrosome markers (Cep135 and γ-tubulin). The fraction of mitotic cells with more than 2 centrosomes was used as a proxy of polyploidy. The frequency of binucleated polyploid cells was evaluated in interphase based on the presence of at least two nuclei and more than two centrosomes per cell. The differential expression of PCNA (S phase) and Ph3 (end of G2/M) markers in the two nuclei of a single cell was scored to evaluate the frequency of asynchrony among the binucleated polyploid cell population.

**Quantitative analysis of DNA damage in single cells – mNSCs**

Z-stacks containing the nuclei of interest were considered in the blue chanel DAPI stained DNA). During the segmentation process, the option of correction to separate the two nuclei of a binucleated mNSC was used if they were considered by the plugin as a unique structure. Quantitative values were obtained using a new plugin developed by QUANTACELL (www.quantacell.com). The FI of the red signal (Ph3) was measured by a thresholding operation. The FI of the green signal (γ-H2Av) was also measured by a thresholding operation and after a number of tests, that took into consideration the γ-H2Ax signal and coverage in terms of Foci considered, we assigned a constant threshold value (210 for measurement of γ-H2Av DNA damage) and multiply the FI obtained by the coverage of the segmented region. The plugin generates the FI of Ph3, which allows the identification of Ph3 (+) and Ph3(-) nuclei in binucleated cells. All the data obtained from individual cell measurements was then plotted in Excel and subsequently in Prism for illustration and statistical analysis. All data plotting and statistical analyses were performed using the GraphPad Prism software.

**Live imaging of *Drosophila* brains**

3$^{rd}$ instar larval brains were dissected in Schneider's *Drosophila* Medium (21720-024, Gibco, TermoFisherScientific, Waltham, MA, USA). The medium was supplemented with 10% heat-inactivated foetal bovine serum (#10500), Penicillin (100 units/ml) and Streptomycin (100μg/ml; Penicillin-Streptomycin #15140), all purchased from Gibco. Brains were placed in medium in a glass bottom 35 mm dish (P35G-1.5-14-C, MatTek Corporation, Ashland, MA, USA). To avoid evaporation and allow gas exchange, brains were covered with a permeable membrane (Standard membrane kit, YSI, Yellow Springs, OH, USA) and the membrane borders were sealed with 10S Voltalef oil (VWR BDH Prolabo, Radnor, PA, USA). For short term live imaging (up to 6 hours), we used 10μl of medium and imaged maximum two different positions. For long term live imaging (> 6 hours), we used 20μl of medium and imaged up to five different positions.

30 Z-stacks of 0.75 to 1μm were acquired at 30 seconds intervals (short term movies) or at 10 minutes intervals (long term movies) on a Yokagawa CSU-X1 spinning head mounted on a Nikon Ti-E inverted microscope, equipped with an EMCCD (electron-multiplying charge-coupled device) 512x512 Evolve camera (Photometrics) using an oil objective 60X NA 1.4. Laser power was kept constant for all the acquisitions. Images were acquired in MetaMorph software and analysed with Fiji. A hyperstack video was created to follow NBs in time and space (x, y and z). All events of interest were manually quantified on raw data.

**Laser ablation**

To verify the efficient recruitment of RpA-70-GFP on sites of DNA damage, 3$^{rd}$ instar larval brains expressing RpA-70-GFP were subjected to laser ablation. We performed short term live imaging using the Yokagawa CSU-X1 spinning head mounted on a Nikon Ti-E inverted microscope described above and a 100X NA 1.3 Oil Iris objective (S Fluor, Nikon). Ablation was carried out on *line scan* regions of interest (1 to 3 ablation pulses of 26 milliseconds, 10 repetitions, laser at maximum power) using an iLas2 system (Roper Scientific France/PICT-IBiSA), piloted by Metamorph (Molecular Devices) and interfaced with a 355 nm UV Q-switched photo ablation laser (50 mW - Output Energy 1.5 μJ; Peak Power 2 kW, Teem Photonics).

**Image processing and statistical analysis**

All images were opened and assembled using Fiji to produce a merged picture or a projection from Z-stacks. They were further processed in Photoshop and image panels were assembled in Illustrator. Statistical analysis was performed using Prism and the tests mentioned in the figure legends. Normal distribution of data was verified by Shapiro-Wilk test.

#### KEY RESOURCES TABLE

| REAGENT or RESOURCE | SOURCE | IDENTIFIER |
|---|---|---|
| Antibodies | | |
| Rabbit anti-aPKC (1:1000) | Santa Cruz | Cat# SC116 |
| Rabbit anti-g-H2aX (1 :500) | Rockland | Histone-H2A pS137 |
| Mouse anti-Ph3 (1 :250) | Sigma-Aldrich | Histone-H3 pS10 |
| Mouse anti-Mab414 | Abcam | ab24609 |
| Rabbit- anti Phospho-histone H3 (pSer10) (1 :750) | Sigma-Aldrich | (H0412) |
| Rabbit- β-catenin (1 :200) | Sigma-Aldrich | C2206 |
| Rabbit- Cep135 (1:500) | Sigma-Aldrich | Generated in the lab |
| Mouse anti-g-H2Ax (1 :750) | Millipore | 05-636 |
| Mouse anti-g-tubulin (1:500) | Sigma-Aldrich | T5326-GTU88 |
| Mouse anti N-cadherin (1:100) | BD Transduction Laboratories | 610921 |
| Mouse anti PCNA (1:500) | Santa Cruz | sc-56 |

| | | |
|---|---|---|
| RFP-Booster (Atto 594) | Chromotek | Cat# rba594; RRID: AB_2631390 |
| ATTO 488-Booster | ChromoTek | AD 488-21 |
| Mouse anti-GFP (clone JL-8) | Clonetech Laboratories | Cat# 632381; RRID: AB_2313808 |
| RFP-Booster (Atto 594) | Chromotek | Cat# rba594; RRID: AB_2631390 |
| 546-conjugated Phalloidin | Thermo Fisher Scientific | A22283 |
| 647-conjugated Phalloidin | Thermo Fisher Scientific | A22287 |
| Goat anti-Rabbit IgG Alexa Fluor 488 (1:250) | Thermo Fisher Scientific | Cat# A-11034, RRID:AB_2576217 |
| Goat anti-Guinea Pig IgG Alexa Fluor 647 (1:250) | Thermo Fisher Scientific | Cat# A-21450, RRID:AB_2735091 |
| Goat anti-Mouse IgG Alexa Fluor 568 (1:250) | Thermo Fisher Scientific | Cat# A-11031, RRID:AB_144696 |
| **Chemicals, Peptides, and Recombinant Proteins** | | |
| 4',6-diamidino-2-phenylindole DAPI | Thermo Fisher Scientific | Cat# 62248 |
| Hoechst | Thermo Fisher Scientific | Cat# 33342 |
| Phosphate-Buffered Saline (PBS) | VWR | Cat# L182-10 |
| RO3360 | Sigma-Aldrich | SML0569 |
| 16% Paraformaldehyde EM Grade | EMS | Cat# 15710 |
| 37% Formaldehyde, Microfiltered | EMS | Cat# 15686 |
| Methanol RPE | CARLO ERBA Reagents | Cat# 414819 |
| Triton X-100 | Euromedex | Cat# 2000-C |
| Sodium Azide | Sigma-Aldrich | Cat# 71289 |
| Schneider's *Drosophila* Medium | Gibco | Cat# 21720024 |
| Fœtal Bovine Serum, heat inactivated | Gibco | Cat# 10500 |
| Penicillin-Streptomycin | Gibco | Cat# 15140 |
| 35 mm glass bottom Dish | MatTek Corporation | Cat# P35G-1.5-14-C |
| Membrane kit, Standard | YSI | Cat# 098094 |

| | | |
|---|---|---|
| Oil 10 S, VOLTALEF® | VWR Chemicals | N/A |
| **Experimental Models: Organisms/Strains** | | |
| W[118] | BDSC | BDSC_5905 |
| WT isogenic | Basto Lab | N/A |
| Actin-GAL4 (act5C-GAL4) | Bloomington Drosophila Stock Center | BDSC_3954 |
| Asense-GAL4 | Lee Lab [15] | N/A |
| $GAL80^{ts}$ ($\alpha$-Tub84B-$GAL80^{ts}$) | BDSC | RRID:BDSC_7017 |
| $\alpha$-Tubulin-GFP | Raff J.W Lab | N/A |
| Histone-RFP (H2Av-mRFP) | BDSC | BDSC_23651 |
| $sqh^1$ / FM7, Kruppel-GAL4,UAS-GFP | Karess R.E Lab | N/A |
| $sqh^1$ ; Tub-GFP,Hist-RFP | This study | N/A |
| $Sqh^{RNAi}$ | Bl#32439 | [7] |
| $Pav^{RNAi}$ | Bl#42573 | [9] |
| $Ani^{RNAi}$ | Bl#42573 | [16] |
| $DSas4^{mut(S2214)}$ | BDSC | BDSC_12119 |
| Tubulin-GFP,Histone-RFP | Basto Lab | N/A |
| Actin-GAL4,$GAL80^{ts}$ | Bellaiche Lab | N/A |
| $sqh^1$ ; Tub-GFP,Hist-RFP | This study | N/A |
| Spc-25-GFP | Lehner C.F. lab | [3] |
| Polo Trap | Raff J.W. lab | [4] |
| RpA-70 GFP | Wiechaus E. lab | [5] |
| Fucci#2 | | [6] |
| **Software and Algorithms** | | |
| Prism 7 | GraphPad software, Inc. | Version 7.0a |

| | | |
|---|---|---|
| Photoshop | Adobe | Version 2017.0.1 |
| Illustrator | Adobe | Version 2017.0.2 |
| Metamorph | Molecular Devices, USA | N/A |
| NIS Element software | Nikon | N/A |
| Fiji | Open Sources | https://fiji.sc/ |
| **Other** | | |
| Cover glasses Menzel gläser, 18x18 mm | VWR | Cat# 631-1331 |
| Cover glasses, square, 22x22 mm | VWR | Cat# 631-0125 |
| Slide Superfrost cut edges | Thermo Fisher Scientific | Cat# AA00008232E00MNT10 |
| Cover glasses, circular, 12 mm ∅ | Marienfeld Superior | Cat# 0111520 |

1. Karess, R.E., Chang, X.J., Edwards, K.A., Kulkarni, S., Aguilera, I., and Kiehart, D.P. (1991). The regulatory light chain of nonmuscle myosin is encoded by spaghetti-squash, a gene required for cytokinesis in Drosophila. Cell 65, 1177-1189.
2. Schuh, M., Lehner, C.F., and Heidmann, S. (2007). Incorporation of Drosophila CID/CENP-A and CENP-C into centromeres during early embryonic anaphase. Curr Biol 17, 237-243.
3. Schittenhelm, R.B., Heeger, S., Althoff, F., Walter, A., Heidmann, S., Mechtler, K., and Lehner, C.F. (2007). Spatial organization of a ubiquitous eukaryotic kinetochore protein network in Drosophila chromosomes. Chromosoma 116, 385-402.
4. Novak, Z.A., Wainman, A., Gartenmann, L., and Raff, J.W. (2016). Cdk1 Phosphorylates Drosophila Sas-4 to Recruit Polo to Daughter Centrioles and Convert Them to Centrosomes. Dev Cell 37, 545-557.
5. Blythe, S.A., and Wieschaus, E.F. (2015). Zygotic genome activation triggers the DNA replication checkpoint at the midblastula transition. Cell 160, 1169-1181.
6. Zielke, N., Korzelius, J., van Straaten, M., Bender, K., Schuhknecht, G.F., Dutta, D., Xiang, J., and Edgar, B.A. (2014). Fly-FUCCI: A versatile tool for studying cell proliferation in complex tissues. Cell Rep 7, 588-598.
7. Weng, M., and Wieschaus, E. (2016). Myosin-dependent remodeling of adherens junctions protects junctions from Snail-dependent disassembly. J Cell Biol 212, 219-229.
8. Ni, J.Q., Zhou, R., Czech, B., Liu, L.P., Holderbaum, L., Yang-Zhou, D., Shim, H.S., Tao, R., Handler, D., Karpowicz, P., et al. (2011). A genome-scale shRNA resource for transgenic RNAi in Drosophila. Nat Methods 8, 405-407.
9. del Castillo, U., Lu, W., Winding, M., Lakonishok, M., and Gelfand, V.I. (2015). Pavarotti/MKLP1 regulates microtubule sliding and neurite outgrowth in Drosophila neurons. Curr Biol 25, 200-205.
10. Rujano, M.A., Sanchez-Pulido, L., Pennetier, C., le Dez, G., and Basto, R. (2013). The microcephaly protein Asp regulates neuroepithelium morphogenesis by controlling the spatial distribution of myosin II. Nat Cell Biol 15, 1294-1306.
11. Casso, D., Ramirez-Weber, F.A., and Kornberg, T.B. (1999). GFP-tagged balancer chromosomes for Drosophila melanogaster. Mech Dev 88, 229-232.
12. Sun, T., Wang, X.J., Xie, S.S., Zhang, D.L., Wang, X.P., Li, B.Q., Ma, W., and Xin, H. (2011). A comparison of proliferative capacity and passaging potential between neural stem and progenitor cells in adherent and neurosphere cultures. Int J Dev Neurosci 29, 723-731.
13. Gogendeau, D., Siudeja, K., Gambarotto, D., Pennetier, C., Bardin, A.J., and Basto, R. (2015). Aneuploidy causes premature differentiation of neural and intestinal stem cells. Nat Commun 6, 8894.
14. Dernburg, A.F., Broman, K.W., Fung, J.C., Marshall, W.F., Philips, J., Agard, D.A., and Sedat, J.W. (1996). Perturbation of nuclear architecture by long-distance chromosome interactions. Cell 85, 745-759.
15. Zhu, S., Lin, S., Kao, C.F., Awasaki, T., Chiang, A.S., and Lee, T. (2006). Gradients of the Drosophila Chinmo BTB-zinc finger protein govern neuronal temporal identity. Cell 127, 409-422.
16. Ni, J.Q., Liu, L.P., Binari, R., Hardy, R., Shim, H.S., Cavallaro, A., Booker, M., Pfeiffer, B.D., Markstein, M., Wang, H., et al. (2009). A Drosophila resource of transgenic RNAi lines for neurogenetics. Genetics 182, 1089-1100.
